# Isolated occurrences of membrane perturbation by mechanosensing from weakly aggregating silver nanoparticles

**DOI:** 10.1101/623678

**Authors:** Marcos Arribas Perez, Oscar H. Moriones, Neus G. Bastús, Victor Puntes, Andrew Nelson, Paul A. Beales

## Abstract

Silver nanoparticles (AgNPs) have wide-ranging applications, including as additives in consumer products and in medical diagnostics and therapy. Therefore understanding how AgNPs interact with biological systems is important for ascertaining any potential health risks due to the likelihood of high levels of human exposure. Besides any severe, acute effects, it is desirable to understand more subtle interactions that could lead to milder, chronic health impacts. Nanoparticles are small enough to be able to enter biological cells and interfere with their internal biochemistry. The initial contact between nanoparticle and cell is at the plasma membrane. To gain fundamental mechanistic insight into AgNP-membrane interactions, we investigate these phenomena in minimal model systems using a wide-range of biophysical approaches applied to lipid vesicles. We find a strong dependence on the medium composition, where colloidally stable AgNPs in a glucose buffer have negligible effect on the membrane. However, at a physiological salt concentrations, the AgNPs start to weakly aggregate and sporadic but significant membrane perturbation events are observed. Under these latter conditions, transient poration and structural remodelling of some vesicle membranes is observed. We observe that the fluidity of giant vesicle membranes universally decreases by an average of 16% across all vesicles. However, we observe a small population of vesicles display a significant change in mechanical properties with lower bending rigidity and higher membrane tension. Therefore we argue that the isolated occurrences of membrane perturbation by AgNPs are due to low probability mechanosensing events of AgNP aggregation at the membrane.

**GRAPHICAL ABSTRACT:** 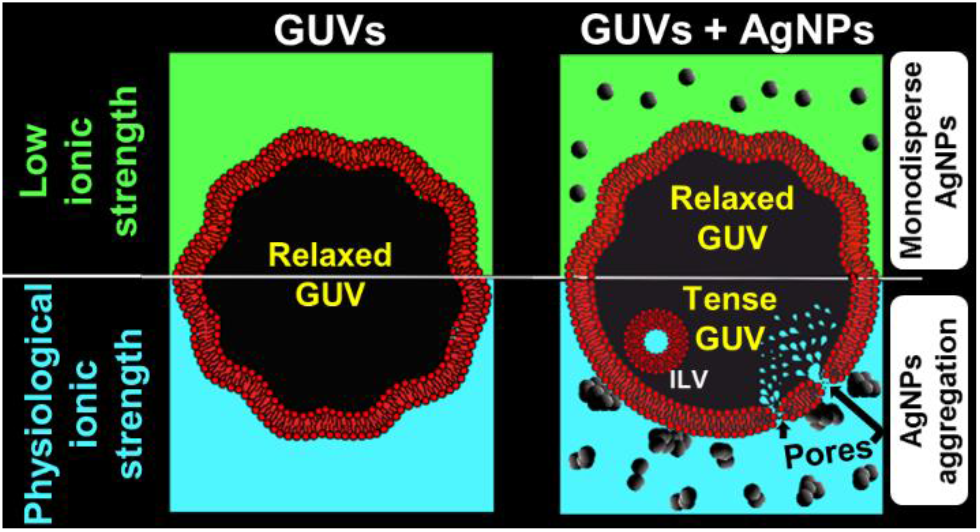

---

Engineered nanomaterials or nanoparticles (NPs) have a very high surface-to-volume ratio that modifies their physicochemical properties due to being largely composed of high energy surface atoms compared to atoms existing in the more stable “bulk”^1^. These novel properties affect the biological activity and biocompatibility of NPs and can lead to advantageous characteristics for their application in biomedicine as therapeutic and diagnosis systems ^2,3^. To perform the desired biomedical function, NPs often must be able to pass across lipid biomembranes to reach specific subcellular targets. However, this NP translocation may result in undesirable membrane perturbations and dysregulation of biochemical processes which can lead to severe cell damage and even cell death ^4–7^.

Noble metal NPs, especially silver nanoparticles (AgNPs), are the most widely used nanomaterials^8,9^. Due to their unique electrical, thermal, optical, antibacterial and anti-inflammatory properties, AgNPs have been largely studied for biomedical applications, such as biosensing, imaging, diagnosis, and antimicrobial therapies^10,11^. Additionally, AgNPs have been proposed as a potential candidate for cancer theranostics, which allows the simultaneous accurate diagnosis and targeted treatment of the disease ^12^. Despite all these advantageous properties, AgNPs have been reported to occasionally cause serious injuries to eukaryotic cells, but the mechanisms behind this cytotoxicity are still not well understood ^13^. In this context, the interaction of AgNPs with the membrane is essential for their biomedical activity but is also the initial step of the toxicity pathway. This interaction often lead to the internalisation of the NPs by the membrane and can induce loss of membrane integrity ^14,15^. Once internalised, AgNPs could cross and damage other sub-cellular membranes to enter important organelles, such as mitochondria, and originate endogenous reactive oxygen species (ROS) ^14–16^. Endogenous ROS are mainly generated by the dysregulation of the respiratory chain of the inner mitochondrial membrane ^17^. While the toxicity of AgNPs is currently primarily attributed to the release of soluble silver ions^18,19^, it is plausible that the increase of endogenous ROS is, at least in part, related with the ability of AgNPs to disrupt lipid membranes ^19^. However, corrosion processes are also REDOX active and induces ROS ^20^, especially in chemically labile NPs, such as AgNPs.

NP-membrane interactions are extremely complex processes and involve several attractive and repulsive forces acting together at the nanoparticle-membrane interface (nanobio interface) ^21^. The nature of the NP-biomembrane interactions and their potential toxicity do not only depend on the composition of the NPs, but are also determined by numerous physicochemical properties of NPs. The size and shape of NPs play a significant role in the interaction mechanism of nanomaterials with biological membranes. Zhang *et al* showed that 18 nm SiO2 NPs cause permanent holes in giant unilamellar vesicles (GUVs) and a decrease in lipid lateral diffusion, whereas SiO2 NPs larger than 78 nm are wrapped by the membrane and lead to an increase in membrane fluidity ^22^. Chithrani *et al* observed higher cellular uptake of spherical gold NPs (AuNPs) into HeLa cells than rod-shaped AuNPs ^23^. Additionally, NP-biomembrane interactions are highly dependent on surface modifications and the charge of NPs. Moghadam *et al* modified the surface coating and charge of AuNPs and titanium dioxide (TiO_2_) NPs and observed that AuNPs and TiO2 NPs with positive charge significantly increase the permeability of DOPC membranes while the dye leakage caused by negatively charged NPs is insignificant ^24^. A similar behaviour has been recently reported by Montis *et al* who showed that cationic AuNPs induce more drastic effects on zwitterionic and negatively charged membranes than anionic and PEG-coated AuNPs ^25^. Moreover, the properties of the surrounding medium, such as pH, temperature, ionic strength, macromolecular crowding, and viscosity can modify many important characteristics of NPs such as surface charge, solubility and colloidal stability, and also can modulate their biological activity ^21,26,27^. The considerably large number of parameters influencing the NP behaviour along with the complexity of biological membranes, makes the understanding of NP-membrane interactions very challenging.

The high complexity of the system represent a limitation for understanding specific mechanisms behind the interaction between NPs and cell membranes. To deal with this issue, it is fundamental to find ways to reconstitute simpler *in vitro* model systems that are easier to control and systematically investigate. One example of such *in vitro* systems are biomimetic model membranes which are synthetic lipid bilayers where the lipid composition can be selected, the lipids can be modified, for example labelled with a fluorescent dye, the number of biomolecular components can be reduced and the medium conditions can be tightly controlled ^28,29^. These artificial model membranes can be investigated using a multitude of biophysical techniques, including calorimetry, spectroscopy and microscopy. This provides detailed information on the structure, mechanics, dynamics and functions of model membranes as well as on their interactions with matter in their local environment, in this case engineered nanoparticles.

In this investigation, we use advanced spectroscopy and microscopy techniques to study changes in the physicochemical properties of lipid membranes upon interaction with AgNPs. Our experiments are performed on large unilamellar vesicles (LUVs) and GUVs. Ensemble characterisation of LUVs (400 nm diameter) provides valuable information about the average behaviour of the vesicles in the sample, whereas GUVs are cell-sized model membranes which are observable by optical microscopy at the single vesicle level and hence allow us to detect rare events, transient processes and study the true distribution of complex behaviours in our experiments ^28^.

## MATERIALS AND METHODS

### Materials

1,2-dioleoyl-sn-glycero-3-phosphocholine (DOPC) lipid and 1,2-dioleoyl-sn-glycero-3-phosphoethanolamine-N-(lissamine rhodamine B sulfonyl) (ammonium salt) (18:1 lissamine rhodamine PE) were purchased from Avanti Polar Lipids Inc. (Alabaster, Alabama, USA). Indium tin oxide (ITO) coated glass slides (surface resistivity 8–12 V sq-1), 4-(2-hydroxyethyl)-1-piperazineethanesulfonic acid (HEPES), silver nitrate (AgNO_3_), trisodium citrate (Na3C6H5O7), and tannic acid (C76H52O46) sodium chloride (NaCl), glucose (C6H12O6), sodium hydroxide (NaOH), and bovine serum albumin (BSA) were obtained from Sigma-Aldrich Co. (Gillingham, UK). 5(6)-carboxyfluorescein, 10 kDa dextran labelled with cascade blue, were purchased from ThermoFisher Scientific Ltd. (Loughborough, Leicestershire, UK). Microscope μ-slide 8 well glass bottom chambers (Ibidi GmbH) were purchased from Thistle Scientific Ltd (Glasgow, UK).

### Buffer composition

The experiments were conducted in two different buffers with high and low ionic strength, respectively. The high ionic strength buffer (HEPES saline buffer) consists of 20 mM HEPES and 150 mM NaCl, resembling the ionic strength of physiological media. In contrast, the low ionic strength buffer (HEPES glucose buffer) also contains 20 mM HEPES but 300 mM glucose instead of salt to maintain the same osmolality of the medium in all the experiments. Both buffers were adjusted to pH 7.4 with NaOH.

### Synthesis of AgNPs

Silver nanocrystals of ~20 nm in diameter were prepared by the seeded-growth method recently reported by Bastús *et al* ^30^. In a typical experiment, 100 mL volume of aqueous solution containing 5 mM of sodium citrate (SC) and 0.1 mM of tannic acid (TA) was prepared and heated up to 100°C with a heating mantle in a three-neck round bottomed flask for 15 minutes under vigorous stirring. A condenser was used to prevent the evaporation of the solvent. After boiling had commenced, 1 mL of 25 mM of AgNO_3_ was injected into this solution. The solution became bright yellow immediately indication the formation of the Ag seeds. Immediately after the synthesis of Ag seeds and in the same vessel, the as-synthetized silver seeds were grown by cooling down the solution to 90°C. First, the seed solution was diluted by extracting 20 mL of sample and adding 17 mL of Milli-Q-water. Then 500 μL of SC [25 mM], 1.5 mL of 2.5 mM TA and 1 mL of 25 mM AgNO_3_ were sequentially injected. This process was repeated up to 2 times, progressively growing the size of the AgNPs until reaching the size ~20 nm (~1.8×10^12^ NPs/mL). The obtained AgNPs were purified by centrifugation and stored in a solution containing both TA and SC.

### UV–Vis Spectroscopy

UV–visible spectra were acquired with a Shimadzu UV-2401 PC spectrophotometer. A 10% (v/v) of AgNP solution was placed in a cell and the spectral analysis was performed in the 300–800 nm wavelength range at room temperature.

### Transmission Electron Microscopy

Transmission electron microscopy (TEM) images were acquired with a FEI Tecnai G2 F20 S-TWIN HR(S) TEM equipped with an energy-dispersive X-ray spectroscopy (EDX) detector, operated at an accelerated voltage of 200 kV. A 10 μL droplet of the sample was drop cast onto a piece of ultrathin carbon-coated 200-mesh copper grid (Ted-pella, Inc.) and left to dry in air. The size of more than 500 particles was computer-analysed using Image J software and measured to calculate size distributions profiles.

### Dynamic light scattering (DLS)

Dynamic light scattering (DLS) and dynamic electrophoretic light scattering (DELSA) were employed to measure the hydrodynamic diameter and the ζ potential of AgNPs, respectively, using a Malvern Zetasizer Nano ZSP (Malvern Panalytical, Malvern, UK) with a 633 nm helium-neon laser. 50 μM AgNPs were suspended in milli-Q water, HEPES saline and HEPES glucose buffer and, after 30 minutes of incubation, each sample was measured three times at a fixed 173° back-scattering angle for the hydrodynamic diameter and 17° scatter angle for ζ potential. The results were processed using the Malvern Zetasizer software to obtain the hydrodynamic diameter from the analysis of the autocorrelation function of the light intensity scattered by the AgNPs, and the ζ potential from the measured electrophoretic mobility using the Smoluchowski approximation. The comparison of the hydrodynamic diameter of AgNPs suspended in the different media was used to evaluate their colloidal stability.

### Preparation of Large Unilamellar Vesicles (LUVs)

Carboxyfluorescein-loaded large unilamellar vesicles (LUVs) were prepared by the extrusion method. Initially, a 25 mg mL^−1^ solution of DOPC in chloroform was dried under high vacuum overnight to get a dry lipid thin film. The lipid film was then rehydrated with 500 μL of 120 mM 5(6)-carboxyfluorescein (CF) solution to form a suspension of liposomes polydisperse in size and lamellarity. Next, to break the multilamellar vesicles (MLVs) and form unilamellar liposomes, the sample was frozen in liquid nitrogen, thawed in water bath at 60°C and vortexed. This freeze-thaw-vortex cycle was carried out 10 times. The sample was subsequently extruded 11 times by passing through a 400 nm pore size polycarbonate membrane (Whatman International Ltd., Maidstone, UK) using an Avanti mini-extruder (Avanti Polar Lipids Inc.) to obtain a homogeneous population of 400 nm LUVs. Finally, the sample was passed through a Sephadex G-25 column to remove the unencapsulated CF via size exclusion chromatography. The size of the LUVs was determined by DLS and the lipid concentration by a standard phosphorus assay ^31^.

### Carboxyfluorescein leakage assay

The leakage assay is a technique that permits the detection of changes in membrane permeability. It is based on the self-quenching ability of the fluorophore CF when it is highly concentrated. The CF was encapsulated within LUVs at enough concentration to be self-quenched and the LUVs were exposed to different concentrations of AgNPs. A membrane damage induced by the AgNPs will produce the release of CF to the external medium where it gets diluted and consequently its fluorescence signal increase significantly. The fluorescence intensity was measured from 500 nm to 600 nm (excitation at 492 nm) using a FluoroMax-Plus spectrofluorometer (Horiba Scientific). The results were calculated from the fluorescence intensity peaks at 514 nm (*I_n_*) and are presented as the normalized percentage of dye leakage. To calculate the fraction of dye release, a sample of LUVs before exposure to AgNPs was used as baseline signal (*I_0_*) which was subtracted from the emission spectrum and values for complete dye leakage (*I_max_*) were obtained by adding 50 μL of the surfactant Triton X-100 which causes the lysis of the LUVs and therefore the release of the 100% of CF. The normalized fraction of CF leakage (*L_n_*) is given by:

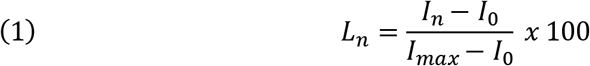

### Preparation of Giant Unilamellar vesicles (GUVs)

GUVs were prepared from 0.7 mM DOPC with 0.5 mol% of 18:1 Lissamine rhodamine PE (Rh-DOPE) dissolved in chlorophorm using the electroformation method ^32,33^. Briefly, 15 μL of lipid solution were deposited as a thin layer over the conductive side of two indium-tin oxide (ITO) coated glass slides and then dried under a nitrogen stream. The ITO slides were then assembled into an electroformation chamber each in contact with a copper tape and separated by a Teflon gasket. The chamber was filled with a 300 mM sucrose solution (300 mOsm kg^−1^) and connected to a function generator to apply an AC electric field. The electroformation was carried out at 5 V peak to peak and 10 Hz for two hours and then the frequency was gradually reduced for approximately 10 minutes to facilitate the closure and detachment of GUVs from the surface. After electroformation, the GUVs were suspended in isotonic (unless otherwise specified) HEPES saline or HEPES glucose buffers. The osmolality of the buffer was measured using a 3320 single-sample micro-osmometer (Advanced Instruments, Norwood, UK). The GUVs observed in this study were between 5 μm and 40 μm (diameter). The GUV-based experiments were conducted at room temperature on a Zeiss LSM 880 inverted laser scanning confocal microscope. The samples were deposited on the microscope slides previously treated with a 10% BSA solution to prevent GUVs from adhering and rupturing onto the glass.

### Determination of changes in morphology and membrane permeability of GUVs

Confocal laser scanning microscopy was used to monitor changes in the permeability and the morphology of GUVs induced by AgNPs. Initially, a membrane-impermeable fluorescent dye (10 kDa dextran cascade blue or CF 576 Da) was added to 200 μL of GUVs suspension, consequently the bulk solution becomes fluorescent whereas the lumen of the GUVs remains uncoloured. The microscope tile scanning option was used to scan large areas of the sample and facilitate the acquisition of statistical data. These scans were acquired before and after incubating the GUVs with different concentrations of AgNPs for 20 ± 5 minutes. The fluorescence intensity in the lumen of GUVs was quantified and normalised using the equation 1 were now *L_n_* is the percentage of dye that leak into a particular GUV, *I_0_* is the average pixel intensity of the dye in the lumen of the GUVs in the unleaked control samples before adding the AgNPs, *I_n_* is the average pixel intensity of the dye in the lumen of the particular GUV and *I_max_* is the average pixel intensity of the dye in the bulk solution. Only the GUVs with a normalised fluorescence intensity in their lumen higher than 20% were considered permeable to the fluorescent molecules in the external medium. The images were analysed using the Fiji extension of ImageJ software (National Institutes of Health, Bethesda, MD) and the proportion of GUVs affected by the AgNPs was determined by manual counting.

### Fluorescence recovery after photobleaching (FRAP)

Fluorescence recovery after photobleaching (FRAP) was used to investigate changes in the fluidity of the membrane. This technique consists of bleaching irreversibly a particular region of interest (ROI) with a high-intensity laser beam and then monitoring, with a low-intensity laser beam, the rate of fluorescence recovery which represents the time needed for the surrounding fluorescent molecules to diffuse into that region of interest ^34^. FRAP experiments were performed on the top pole of GUVs before and after 20 ± 5 minutes of exposure to AgNPs. A circular ROI of 5 ± 0.5 μm diameter was exposed to 5 bleaching scans at 100% laser power and the recovery was monitored by recording time series of 100 frames with the confocal pinhole adjusted to 3 μm. The recovery curves were fitted with Origin Pro using the classic fluorescence recovery model ^35,36^:

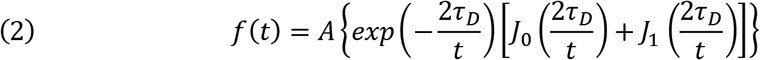

where *t* is time, A is the recovery level, *τ_D_* is the half recovery time, and *J_0_* and *J_1_* are modified Bessel functions of the first kind. The diffusion coefficient (*D*) can then be calculated from the recovery times and the radius of the bleached region (*r*) using:

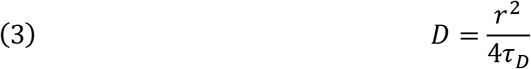

### Flicker spectroscopy

The membrane mechanical properties of GUVs were determined using flicker spectroscopy ^37–39^. This is a non-invasive image analysis technique which quantifies the amplitude of membrane thermal fluctuations (〈|*u(q)*|^2^〉) as a function of their wavenumber (*q=2π/l*) along the length (*l*) of the GUV contour. For these experiments, an osmotic relaxation of the GUVs was carried out incubating the GUVs overnight at 4° C with a hyperosmotic buffer (315 mOsm kg^−1^). Confocal microscopy time series of 1000 frames and a resolution of 1024 × 1024 pixels were taken at the equatorial plane of single GUVs before and after 20 ± 5 minutes of exposure to AgNPs. The confocal pinhole aperture was adjusted to 0.7 μm and, to increase the scan speed, single GUVs were zoomed in upon to the maximum magnification that allows imaging of the whole vesicle. The data was analysed using MATLAB contour analysis software kindly provided by Prof. Pietro Cicuta and co-workers at the University of Cambridge, UK. This programme analyses each frame of the time series and quantifies the membrane tension (*σ*) and bending rigidity modulus (*κ_b_*) by fitting the fluctuation spectrum with the following equation ^39^:

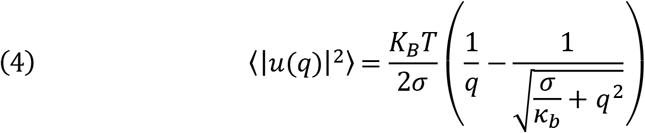

## RESULTS

### AgNPs tend to aggregate in physiological ionic strength buffer

The AgNPs employed in this investigation were spherical with a diameter of 22.4 ± 2.51 nm as determined by transmission electron microscopy (TEM) (Figure 1a). Nonetheless, the conditions of the medium can have an important influence on the biological activity of AgNPs by modifying their physicochemical properties leading, for instance, to nanoparticle dissolution, nanoparticle aggregation or interaction with organic matter ^40^.

**Figure 1.**
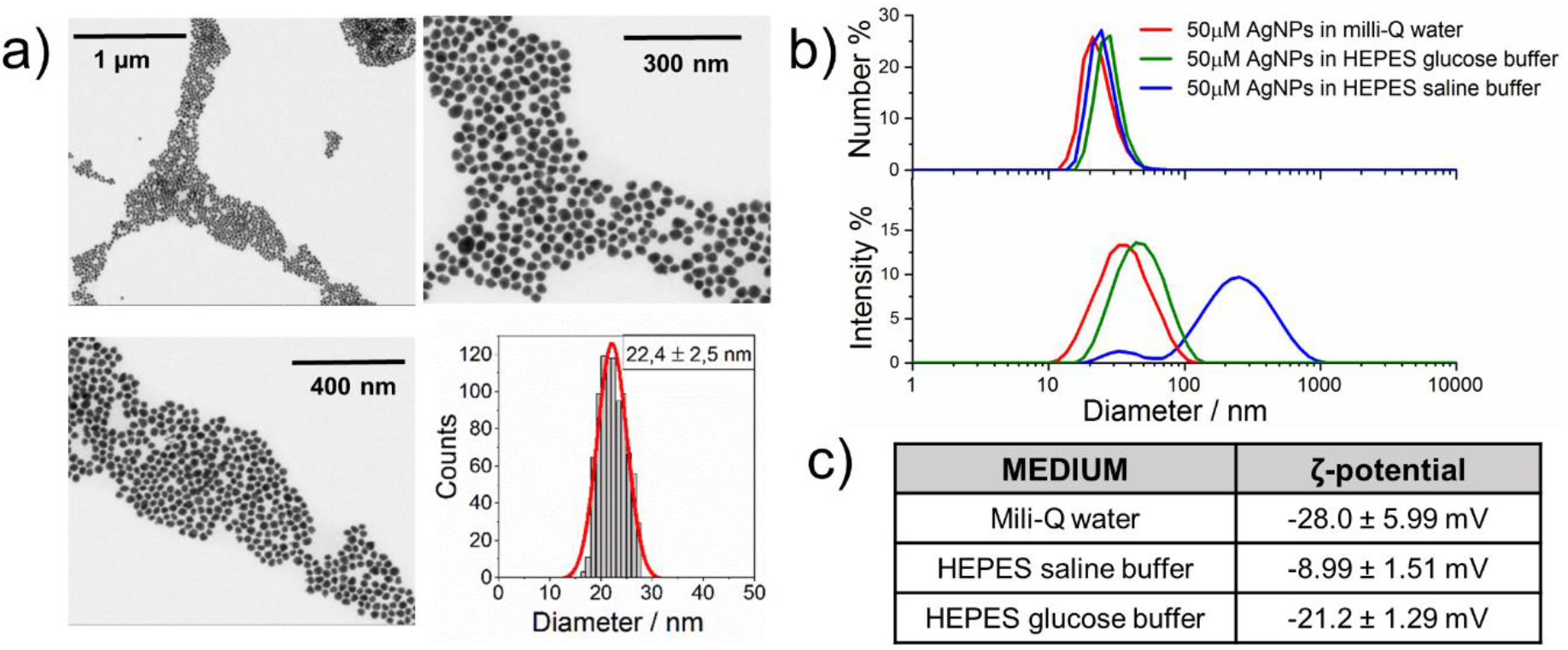
Characterisation of AgNPs in different medium conditions. **a)** TEM images show spherical AgNPs with an average diameter of 22.4 ± 2.51 nm **b)** Dynamic light scattering (DLS) data of 50 μM AgNPs suspended in mili-Q water, HEPES saline and HEPES glucose buffer for 30 minutes. The size distribution by number (top graph) shows a monodisperse distribution of AgNPs with peaks around 25 nm, however the size distribution by intensity (bottom graph) displays similar hydrodynamic diameters of AgNPs dispersed in mili-Q water (40.25 ± 1.02 nm) and HEPES glucose buffer (43.55 ± 0.86 nm), whereas in HEPES saline buffer AgNPs show a tendency to form aggregates (199.40 ± 21.45 nm). c) The ζ-potential of AgNPs dispersed in mili-Q water and HEPES glucose was also similar while in HEPES saline buffer the surface charge of AgNPs becomes less negative.

The colloidal stability and ζ-potential of AgNPs in physiological and low ionic strength buffers was tested using dynamic light scattering (DLS) and electrophoretic light scattering (DELSA). The aggregation of NPs can be easily detected by comparing the hydrodynamic size of AgNPs suspended in milli-Q water with their hydrodynamic size when suspended in the buffer of interest. The size distribution by intensity shows that the hydrodynamic diameter of AgNPs in HEPES glucose buffer (43.55 ± 0.86 nm) does not differs significantly from the results in milli-Q water (40.25 ± 1.02 nm), hence the NPs are stable in this buffer (Figure 1b). The larger size obtained by DLS compared to TEM is expected because while TEM measures only the physical size of the core NPs, DLS measures the hydrodynamic size which in addition to the NP core, takes into account the stabiliser coating and the electrical double layer around the NPs^41,42^. On the contrary, these AgNPs show a tendency to form aggregates (199.40 ± 21.45 nm) when the ionic strength of the medium is high (physiological), however the size distribution by number indicates that at this concentration (50 μM) and incubation time (30 min) most of the AgNPs are still monodisperse (Figure 1b). Similarly, the ζ-potential of AgNPs is comparable in milli-Q water and HEPES glucose buffer, −28.0 ± 5.99 mV and −21.2 ± 1.29 mV respectively, whereas in HEPES saline buffer the surface charge of AgNPs becomes less negative (−8.99 ± 1.51 mV) (Figure 1c). The aggregation tendency and the less negative surface charge of AgNPs in physiological ionic strength conditions are closely related. The negative surface charge provided by the citrate coating stabilises the AgNPs suspension by electrostatic repulsions, however in HEPES saline buffer the high concentration of ions produces a screening of the surface charge that compresses the electrical double layer around the AgNPs^43–45^. Consequently, the electrostatic repulsions between AgNPs become weaker and their aggregation tendency increases.

### The ionic strength of the medium modulates the effect of AgNPs on the membrane permeability

The effect of AgNPs on the membrane permeability was initially investigated by quantifying the release of 5(6)-carboxyfluorescein from DOPC LUVs. The carboxyfluorescein (CF) loaded DOPC LUVs (400 nm diameter) were suspended in isotonic HEPES saline buffer or HEPES glucose buffer reaching a final phospholipid concentration of 0.10 ± 0.02 μM. The LUV suspensions were then exposed to various concentrations of AgNPs (1 μM, 3 μM, 10 μM, 30 μM and 100 μM) and the fluorescence emission of each sample at 514 nm was measured every 15 minutes during a total time of 90 minutes (Figure 2a). The maximum absorption peak of these AgNPs was at 417.4 nm (Figure S1), far from the working wavelength of CF, however this peak can change due to NP aggregation. Thus, control experiments were performed comparing the fluorescence emission of samples at various concentrations of CF and the same samples plus 100 μM AgNPs to ensure that the NPs are not interfering with the fluorescence signal of CF (Figure S2). In HEPES saline buffer, the exposure of LUVs to 10 μM AgNPs produce a notable dye leakage, but at higher AgNPs concentrations dye release from the LUVs begin to be more extensive, reaching a maximum leakage of nearly 15% and 30% after 30 minutes of incubation with 30 μM and 100 μM AgNPs, respectively. At that moment, the dye release reaches a plateau and remains stable for the next 60 minutes. In HEPES glucose buffer, after 30 minutes incubation at the highest concentrations of AgNPs (30 μM and 100 μM), the LUVs produce just a marginal dye release, which does not vary until 90 minutes of exposure when it increases slightly but does not exceed 10% CF release. Since aggregation is known to be concentration and time dependent, at lower concentrations the NPs are less aggregated. However, at the same incubation times, the highest concentrations of NPs lead to more and larger aggregates, but still in suspension, which would be able to interact with the vesicles and perturb their membrane due to their reduced solubility. At longer times, the aggregates would be expected to grow further until they reach a size where they are no longer colloidal, drop out of suspension and sediment to the bottom of the sample. In this way they could become excluded from the solution and no longer able to interact with the vesicles. These results indicate that the conditions of the surrounding environment have a pronounced influence on the interaction mechanism between AgNPs and zwitterionic phospholipid membranes.

**Figure 2.**
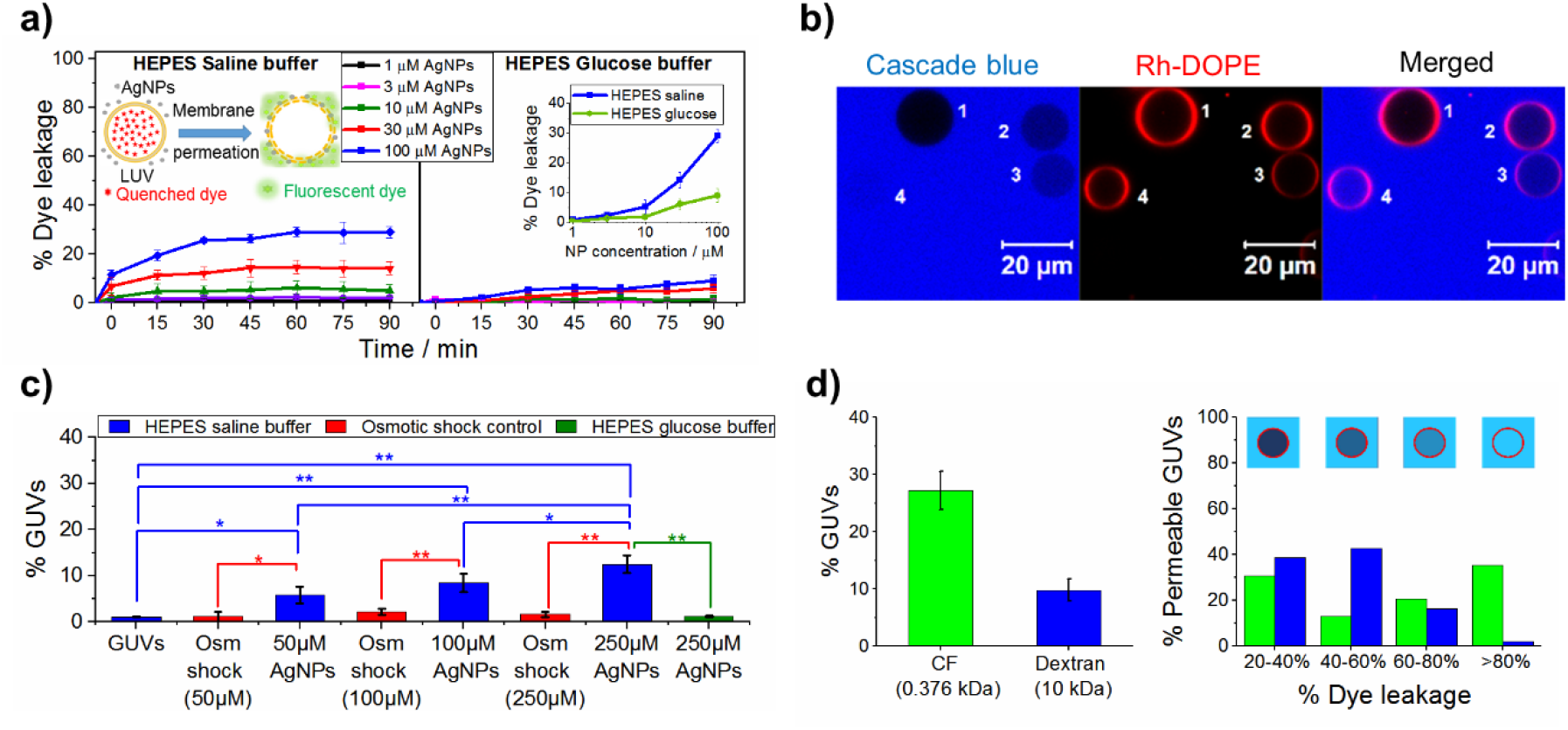
AgNPs in physiological ionic conditions have a more significant impact on membrane permeability. **a)** In HEPES saline buffer AgNPs induce a higher dose-dependent leakage of 5(6)-carboxyfluorescein (CF) from DOPC LUVs than in HEPES glucose buffer. Insets show a schematic representation of CF leakage from LUVs as well as the proportion of dye leakage as a function of NP concentration after 90 minutes of incubation in both buffers. **b)** DOPC GUVs labelled with Rh-DOPE with different degrees of permeability to 10 kDa dextran labelled with cascade blue fluorophore: unleaked GUV (1), partially leaked GUVs (2 and 3) and fully leaked GUV (4). c) The proportion of DOPC GUVs permeable to 10 kDa dextran increases with the concentration of AgNPs in physiological ionic strength medium. Osmotic shock controls were performed to ensure that this effect was not induced by changes in the osmolarity of the medium. In HEPES glucose buffer, the highest concentration of AgNPs does not induce a noticeable change in the number of GUVs permeable to dextran. The statistical significance was tested using a one way ANOVA with Bonferroni multiple comparisons test (*p ≤ 0.05, **p ≤ 0.01). d) Comparison between influx of CF and 10 kDa dextran into DOPC GUVs after exposure to 100 μM AgNPs. A higher proportion of GUVs become permeable to CF than to dextran. The distribution of permeable GUVs according to their level of leakage show a higher influx rate of CF than of dextran. Inset: schematic view of different levels of dye leakage.

The release of the dye encapsulated in the vesicles can be a consequence of different processes, from complete lysis of liposomes to the formation of nanosized pores in the membrane. To identify the mechanism behind the change in membrane permeability, we used confocal microscopy to directly observe the influx of membrane impermeable dextran molecules (10 kDa) labelled with the fluorophore Cascade blue into GUVs after incubating them for 20 ± 5 minutes with 50 μM, 100 μM and 250 μM AgNPs (Figure 2b). Additionally, control experiments were carried out adding equivalent volumes of milli-Q water to the GUVs to test the effect of a potential osmotic shock and ensure that the effects observed are indeed produced by the AgNPs. At physiological ionic strength, the exposure to AgNPs induces a dose-dependent increase in GUVs permeable to 10 kDa dextran, which is statistically significant at the three concentrations of AgNPs tested (Figure 2c). This effect is not observed in low ionic strength (HEPES glucose buffer) conditions where the exposure of the GUVs to AgNPs barely produces any change in their permeability to dextran 10 kDa. Thus, these observations demonstrate a regulating role of the medium composition on the AgNPs-membrane interactions.

The influx of large macromolecules, such as 10 kDa dextran, into the lumen of GUVs must be induced by the formation of pores in the membrane, which can vary in size and lifetime ^46–48^. Confocal microscopy enables the analysis of the behaviour of individual GUVs in the sample, allowing us to quantify the specific degree of leakage in each individual vesicle observed (Figure 2b). Figure 2d compares the proportion of GUVs permeable to CF (0.37 kDa) and 10 kDa dextran after exposure to 100 μM of AgNPs as well as the distribution according to their degree of leakage. The proportion of GUVs permeable to CF molecules is nearly three times higher than to macromolecules of dextran. The distribution of permeable GUVs as a function of the normalised fluorescence intensity in their lumen show that nearly 40% of the permeable GUVs were fully filled (> 80% dye leakage) with CF after exposure to 100 μM AgNPs whereas less than the 20% of GUVs permeable to 10 kDa dextran showed more than 60% of leakage, of which just a marginal proportion were fully leaked. According to these results, the AgNPs produce nanoscale pores in physiological ionic solutions, which allow the easier and faster transmembrane diffusion of small molecules than larger macromolecules. Moreover, the fact that just a low proportion of the GUVs get fully leaked suggest that the pores formed are transient and the membrane can self-heal, recovering its integrity and blocking further transmembrane diffusion.

### AgNPs can induce the formation of intraluminal vesicles in physiological ionic strength conditions

The exposure to AgNPs produces membrane intraluminal vesicles (ILVs) in a small proportion of the GUVs when suspended in physiological ionic strength buffer. These ILVs are small vesicles filled with extravesicular bulk medium (10 kDa dextran labelled with cascade blue fluorophore) within the lumen of the GUVs. The formation of the ILVs is very fast, however some images show potential intermediate states consisting of pearling tubes (Figure 3). This phenomenon has been previously reported by Yu and Granick, who observed that aliphatic amine NPs encapsulated within DOPC GUVs adsorb onto the inner lipid leaflet of the membrane and induce an initial protrusion of large tubes followed by pearling events^49^. Montis *et al*. also observed that the exposure of POPC GUVs to cationic gold nanorods (AuNR) produce tubular lipid protrusions that breakup into pearls^25^. The experimental procedure and data analysis used for these experiments was the same employed for the estimation of dye leakage into GUVs. The proportion of GUVs with ILVs is low but statistically significant with respect to GUVs before exposure to AgNPs and the osmotic shock controls. Notably, ILVs formation does not vary significantly with the concentration of AgNPs, presenting in all cases between the 5% and 6% of GUVs observed (Figure 3). This effect was not seen in low ionic strength buffer, hence it is also influenced by the composition of the medium.

**Figure 3.**
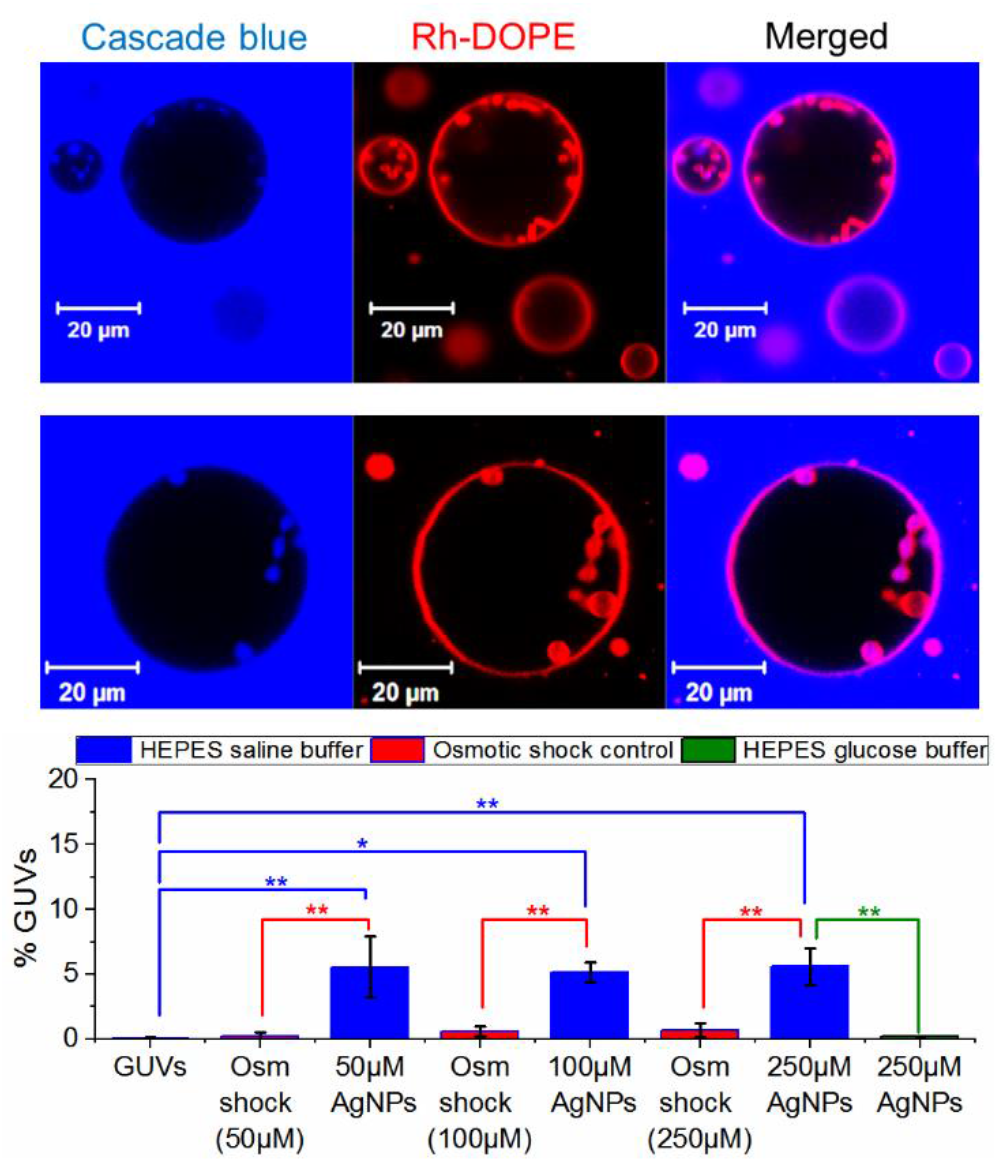
AgNPs in physiological ionic conditions can induce topological changes in GUV membranes with low probability. Small intraluminal vesicles (ILVs) and tubular structures filled with bulk solution are observed inside the GUVs upon exposure to AgNPs in physiologic ionic strength conditions. The proportion of GUVs with ILVs observed is low and similar at the three concentrations of AgNPs explored. Osmotic shock controls were performed to ensure that this effect was not induced by changes in the osmolarity of the medium. In HEPES Glucose buffer, the exposure to 250 μM of AgNPs do not favour the formation of membrane inclusions. The statistical significance was tested using a one way ANOVA with Bonferroni multiple comparisons test (*p ≤ 0.05, **p ≤ 0.01).

These results suggest that the formation of ILVs is a stochastic event that has plateaued at its maximal extent by 50 μM AgNPs. We propose that this effect could be related to the aggregation of AgNPs in physiological conditions. Previous work on ZnO NPs reported that largescale aggregation leading to microscale aggregates that drop out of suspension have reduced membrane interactions^50^. The nanoscale aggregates we observe for AgNPs suggest they are more weakly aggregating and maintain their solution dispersion, this time increasing their activity at membranes. The aggregation of NPs can be considered as a stochastic process which may lead to a large number of aggregates polydisperse in size and shape. Computer simulation studies have shown that the configuration that NPs adopt to form clusters or aggregates modifies their ability to bend lipid membranes ^51–54^. Therefore, the formation of ILVs could be a result from the assembly of AgNPs clusters on the GUV membrane. Nonetheless, only aggregates with a particular shape, size and orientation would be able to efficiently bend the membrane to induce pearling and ILV formation.

### AgNPs slightly modifies the membrane fluidity in physiological ionic strength conditions

Fluorescence recovery after photobleaching (FRAP) was employed to investigate changes in lipid lateral mobility. The diffusion coefficients were calculated from the mobility of the fluorescent lipids (Rh-DOPE) within the membrane of GUVs estimated from the fluorescence recovery times in a previously bleached region of the upper pole of a vesicle. The diffusion coefficient of DOPC GUVs was calculated before and after incubation with AgNPs in both high and low ionic strength buffers. Between 15 and 20 GUVs were analysed for each condition and the results are summarised in Figure 4, which show the distribution of lipid diffusion coefficients in GUVs as well as an example of FRAP recovery curve in the two media. In HEPES saline buffer, the diffusion coefficient of the DOPC lipids after exposure to 100 μM AgNPs drops slightly from 3.02 ± 0.34 μm^2^ s^−1^ to 2.54 ± 0.31 μm^2^ s^−1^. We suggest that this subtle impact of AgNPs on the fluidity of the membrane is originated by a slight increase in the lipid packing produced by generic adsorption interactions of AgNPs on the membrane. Notably, this decrease in membrane fluidity occurs universally across all GUVs in the sample as the histograms are almost fully displaced from one another in the physiological buffer, unlike the low probability effects of poration and ILV formation seen in earlier experiments. This implies that the change in membrane fluidity in itself is not the primary mechanism that gives rise to these other membrane perturbations.

**Figure 4.**
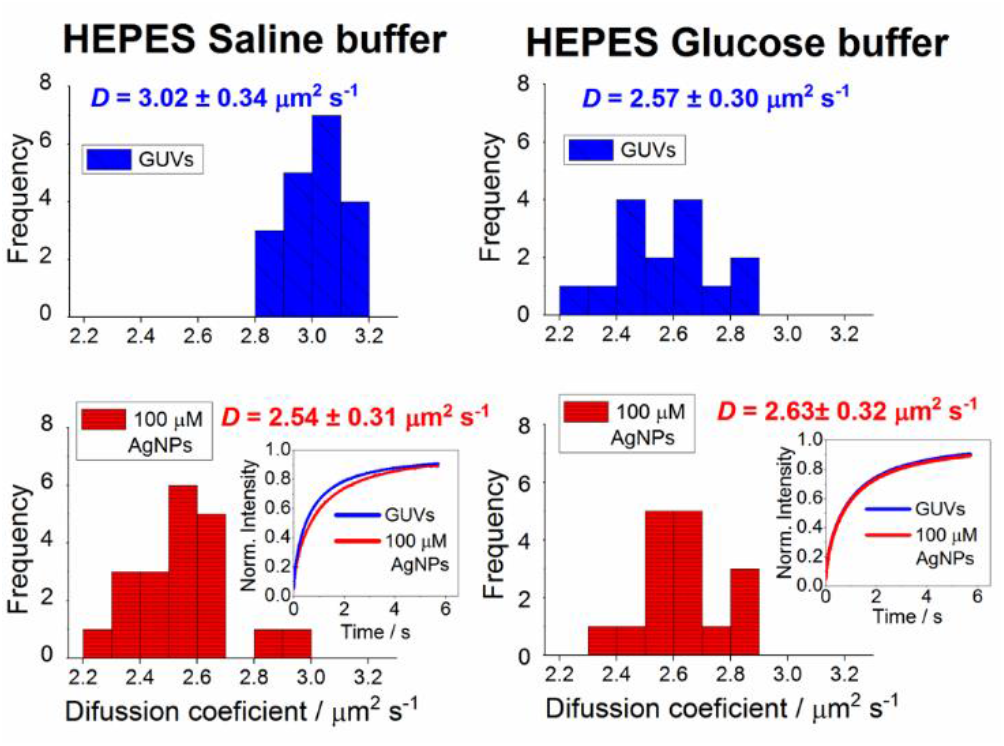
AgNPs in physiological salt conditions decrease membrane fluidity but have no impact in low ionic strength conditions. Distribution of diffusion coefficients obtained from FRAP recovery curves of DOPC GUVs in HEPES saline buffer (high ionic strength) and HEPES glucose buffer (low ionic strength) before and after exposure to 100 μM AgNPs. *D* values indicate the mean diffusion coefficient calculated in each condition. In HEPES saline buffer, AgNPs induce a slight but statistically significant decrease in lipid lateral diffusion. In low ionic strength medium the membrane fluidity of DOPC GUVs is lower than in HEPES saline and does not change after incubation with AgNPs. Insets show examples of FRAP recovery curves obtained in each condition.

In low ionic strength conditions, the AgNPs barely modify the lipid lateral mobility suggesting negligible or very transient adsorption of AgNPs onto the membrane under these conditions. Interestingly, the average diffusion coefficient of DOPC GUVs in these conditions (2.54 ± 0.31 μm^2^ s^−1^) is lower than in high ionic strength buffer. These data suggest that the ionic strength of the medium could not be the only environmental condition modulating the interaction mechanism between the AgNPs and the membrane, but also the presence of high concentrations of sugars might be protecting the membrane. Previous studies have reported that sugars decrease the lipid lateral diffusion in a concentration-dependent manner using Fluorescence Correlation Spectroscopy (FCS) ^55^. Therefore, we attributed the lower membrane fluidity in low ionic strength buffer to the high concentration of glucose in the medium.

### AgNPs considerably affect the mechanical properties of a sub-population of GUVs in physiological ionic strength buffer

The ability of AgNPs to produce membrane invaginations suggest that these NPs might modify the mechanical properties of the membrane. Flicker spectroscopy experiments were performed to quantify the distribution of membrane tension (*σ*) and bending modulus (*K_b_*) of GUV membranes. This technique has a limitation in spatiotemporal resolution which does not allow us to calculate accurately the amplitude of membrane fluctuations (〈|*u(q)*|^2^)) of high-tension GUVs within a broad enough range of wavenumbers (*q*)^56^. This issue was circumvented by the osmotic relaxation of GUVs which decreases the tension of the membrane, increasing the amplitude of the membrane’s thermal undulations and therefore making the fitting of the power spectrum more reliable. Additionally, the osmotic pressure of the AgNPs suspension was balanced with sucrose until isotonic to the experimental medium to prevent changes in the osmolarity of the medium during the experiment.

As expected, in low ionic strength glucose buffer AgNPs do not induce any significant change in either the tension or the bending rigidity of the membrane. In high ionic strength (physiological) conditions, the addition of 100 μM AgNPs to the osmotically relaxed GUVs induces subtle changes in the mechanical properties of the membrane. The distribution of both the tension and the bending rigidity values become wider after the incubation of GUVs with AgNPs. We observe a rise in the mean membrane tension, which despite not seeming to be a drastic change, is statistically significant (p < 0.01) (Figure 5a). This increase in membrane tension is accompanied by a small decrease in membrane rigidity, nevertheless this latter change does not show statistical significance (Figure 5b). Interestingly, these changes in membrane mechanics are not produced by a global effect on every GUV in the sample, but arise from profound changes in membrane tension and bending rigidity of just a fraction of the GUVs analysed (Figure 5c). These results could be directly related to the formation of membrane pores, invaginations and ILVs, described early, which also only occur in a small sub-population of the GUVs. A recent study has shown that formation of ILVs by the endosomal sorting complex required for transport (ESCRT) produces a significant increase in the membrane tension of GUVs originated by the removal of excess membrane surface area ^57^. Furthermore, the increase of the membrane tension is known to favour the membrane poration^58–60^, and therefore could favour the membrane permeation effect induced by AgNPs.

**Figure 5.**
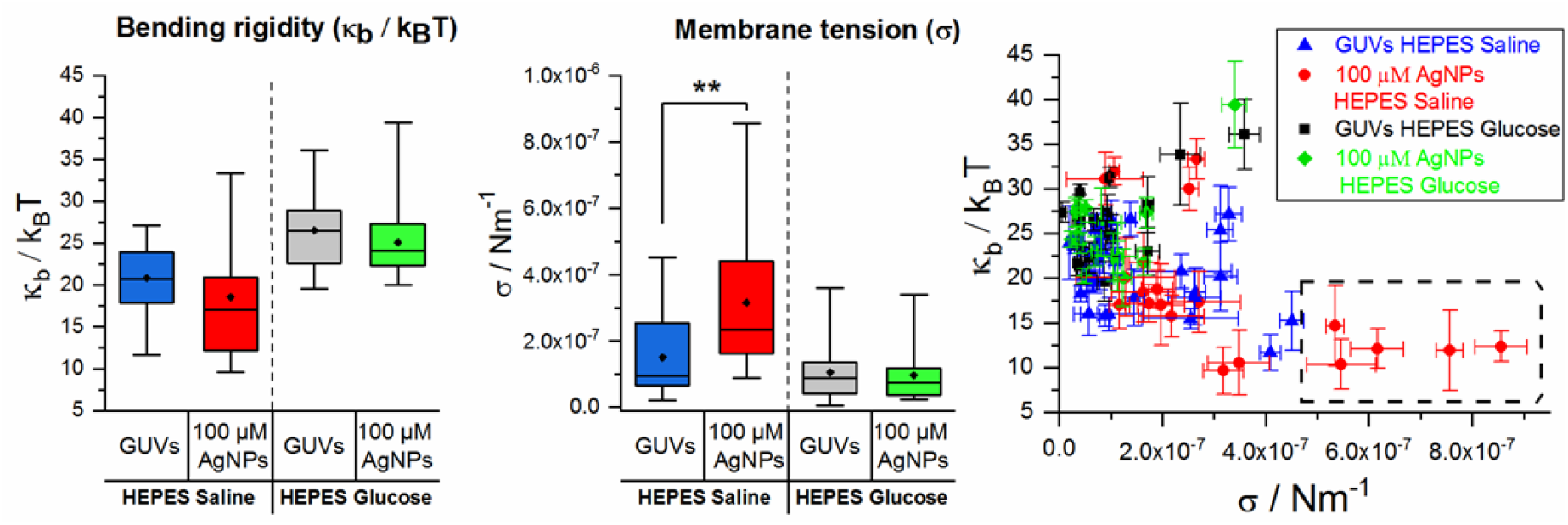
AgNPs in physiological ionic conditions significantly impact the membrane mechanical properties of a sub-population of GUVs. Effect of AgNPs on the mechanical properties of osmotically relaxed DOPC GUVs. **a)** In HEPES saline, 100 μM AgNPs induce a broader distribution and a slight shift in bending rigidity (*κ_b_*/*k_B_T*) which is not statistically significant. In low ionic strength buffer the bending rigidity barely changes. **b)** In HEPES glucose buffer the membrane tension (σ) does not vary after exposure of GUVs to 100 μM AgNPs whereas is HEPES saline a significant change in membrane tension is observed (**p ≤ 0.01, ANOVA with Bonferroni multiple comparisons test). **c)** The plot of bending rigidity against membrane tension show that the majority of GUVs in all conditions have similar membrane tension (>5×10^−7^ Nm^−1^) and bending rigidity (15 – 35 *κ_b_* / *k_B_T*) However, a small proportion of GUVs in HEPES saline experience a great change in membrane tension after exposure to 100 μM AgNPs which is also associated to a small decrease in bending rigidity (dashed box). These GUVs show a membrane tension higher than 5×10^−7^ Nm^−1^ and a bending rigidity between 10 and 15 *κ_b_* / *k_B_T*.

## DISCUSSION

The behaviour of nanomaterials in biological systems is governed by their physicochemical properties, nonetheless these properties are susceptible to change once the NPs enter biological media. The results presented in this work firstly evidence a significant impact of the medium conditions on the surface charge and colloidal stability of AgNPs. The citrate coating of AgNPs gives them a negative surface charge and stabilises the colloidal suspension in low ionic strength media through electrostatic double-layer repulsions. However, the high concentration of ions in physiological conditions modifies the ζ-potential of AgNPs and promotes their aggregation making the NPs less negatively charged. In this medium, Na+ ions screen the negative surface charge and decreases the inter-particle repulsive force, thereby facilitating the concentration dependent aggregation of AgNPs ^61,62^.

Several previous investigations have focused on the effect of the attachment of proteins to the NP surface (protein corona) on the biological activities of NPs ^50,63–68^, however comparatively less attention has been paid to other properties of the medium, such as pH or ionic strength, which can also alter the physicochemical properties of NPs and their biointeractions. Our data show that in high (physiological) ionic strength conditions, AgNPs induce subtle but important effects on the physicochemical properties of the membrane, whereas in low ionic strength buffer the membrane maintains its integrity after exposure to AgNPs. Therefore, the buffer conditions modify the interactions of AgNPs acting at the nanobio interface and hence modulate their interaction with membranes. At pH 7.4, the DOPC membrane carries a slight negative charge ^69^. Thus, in a low ionic strength environments, electrostatic repulsion between AgNPs and the DOPC bilayer may dominate over other attractive forces, preventing significant interaction. However, when the ionic strength increases, the screening of surface charge of AgNPs decrease electrostatic repulsive forces between the NPs and the membrane, increasing the likelihood of more significant interaction. In addition, the loss of colloidal stability and weak aggregation behaviour of AgNPs makes them less soluble and more surface active, further increasing membrane interactions. This modulatory effect of the ionic strength of the medium on the AgNPs-membrane interaction has been reported earlier by Wang *et al*, who, using quartz crystal microbalance with dissipation (QCM-D), found that the deposition rates of AgNPs on DOPC supported lipid bilayers (SLBs) increases when the concentration of salt in the medium rises ^70^. A similar electrostatically-mediated interaction was reported by Li and Malmstadt for cationic polystyrene NPs (PNPs) which interact weakly with the membrane as the ionic strength of the medium increases ^46^.

In physiological conditions, we observe that AgNPs induce changes in membrane permeability and can form membrane invaginations such as ILVs. Several studies have shown the ability of eukaryotic, bacterial and virus proteins as well as antibacterial peptides and NPs to change the morphology of the membrane and form invaginations and ILVs without the need of cellular endocytic mechanisms or external sources of energy ^25,49,57,71–74^. Generally the interaction of single proteins or particles is not strong enough to induce these large deformations of the membrane and thus many molecules or particles must cooperate to bend the membrane ^51,72^. For instance, the clustering of Gb3-binding B subunit of the bacterial Shiga toxin (STxB) can induce membrane invagination in artificial model membranes, however these invaginations are not observed when the clustering is inhibited ^71^. We propose that single AgNPs do not possess enough energy to bend the membrane, nonetheless the formation of NP clusters of a particular size and shape will increase their ability to deform the membrane and produce invaginations and ILVs. The formation of these structures removes excess membrane, which, along with the pressure generated by the AgNPs that adhere onto the membrane, increases the membrane tension and can lead to membrane poration ^46^. The increase in membrane tension is also known to be the driving force of opening of pores which will in turn produce membrane permeation, causing the membrane translocation of impermeable dextran probes and the relaxation of the membrane tension. The lifetime of membrane pores is usually short because as the membrane tension relaxes, the line tension at the pore edge drives closure of the pore ^58,75^. The size of the pore defines the minimum size of the molecules that can diffuse across the membrane. The higher permeability to carboxyfluorescein (0.37 kDa) than to dextran (10 kDa) observed in our experiments represents the presence of nanoscale pores in the membrane as a result of the AgNPs. Furthermore, the fact that most of the GUVs observed were not fully leaked indicates these pores are transient.

In general, toxicology studies focus on terminal effects where the analytes induce severe damages in the membrane or other cellular components that lead to cell death. However, other subtle effects in the physicochemical properties of the membrane can also have biological importance. Cells are able to sense mechanical stimuli and convert them into intracellular biochemical signals to adapt to their microenvironment ^76^. Mechanical forces can modify the mechanical and dynamical properties of the membrane, which in turn can induce conformational changes in membrane proteins, such as ion channels^77^, G-protein coupled receptors (GPCRs) ^78–80^ and integrins^81^. These proteins trigger metabolic cascades that lead to different cellular responses such as cell migration, differentiation, and proliferation^82,83^. In endothelial cells for instance, the plasma membrane senses haemodynamical forces generated by the blood activating downstream signalling pathways related with inflammatory responses, regulation of blood pressure or coagulation processes ^84^. Another important example of mechanical sensing and transduction is the Hippo pathway, which controls organ growth by regulating cell proliferation ^85^. The mechanical stress applied to the plasma membrane modulates the actin cytoskeleton and activate GPCRs. This begins a complex signal pathway that eventually activates the proto-oncogenes proteins YAP/TAZ which translocate from the cytoplasm to the nucleus and induce cell proliferation ^86^. Prolonged mechanical stress can lead to an overexpression of YAP/TAZ promoting unregulated cell proliferation and eventually oncogenesis ^85,86^. Therefore, even small changes in the mechanical and dynamical properties of the membrane, such as the ones produced by AgNPs, can induce multiple cellular responses which lead to a myriad of processes encompassing from inflammatory responses to the development of serious diseases.

Biological fluids are complex and crowded with biomolecules which can interact with the NPs in a non-specific manner and modulate their behaviour. Thereby, the biological interactions of NPs can be influenced by multiple external factors. These factors can be simulated and controlled *in vitro*, for instance introducing proteins to investigate the formation of protein coronas and their effect in NP-membrane interactions. Another factor that would be interesting to study is the impact of excluded volume effects generated by macromolecular crowding on these interactions. This phenomenon is known to promote protein-protein interactions ^87^, and the binding of proteins to the membrane ^88^, thus it is likely to influence the interaction between NPs and the membrane.

To summarise, here we show the significant impact that the ionic strength of the medium have on the physicochemical properties of AgNPs and their interactions with biomembranes. From our results we propose that monodisperse AgNPs are non-interacting and could be safely exploited as imaging contrast agents for diagnosis and cancer theranostics. However, the aggregation of AgNPs would lead to two different possibilities: i) large aggregates precipitate and are expelled from the solution and do not interact with the membrane. These large aggregates are expected to be highly toxic because they are more difficult to be transported in the blood and can be accumulated in certain organs before reaching their specific targets and produce severe damage ^20^. ii) Smaller aggregates become more membrane active than monodisperse AgNPs increasing the tension of the membrane and opening pores. The formation of transient pores offers opportunities in transfection technologies but at the same time raises nanotoxicology concerns.

## ACKNOWLEDGEMENTS

This work was funded by the E.U. Horizon 2020 project HISENTS (GA No. 685817). MAP acknowledges funding from an endowed PhD scholarship at the University of Leeds.

## SUPPLEMENTAL INFORMATION

### UV–Vis Spectroscopy

**Figure S1.**
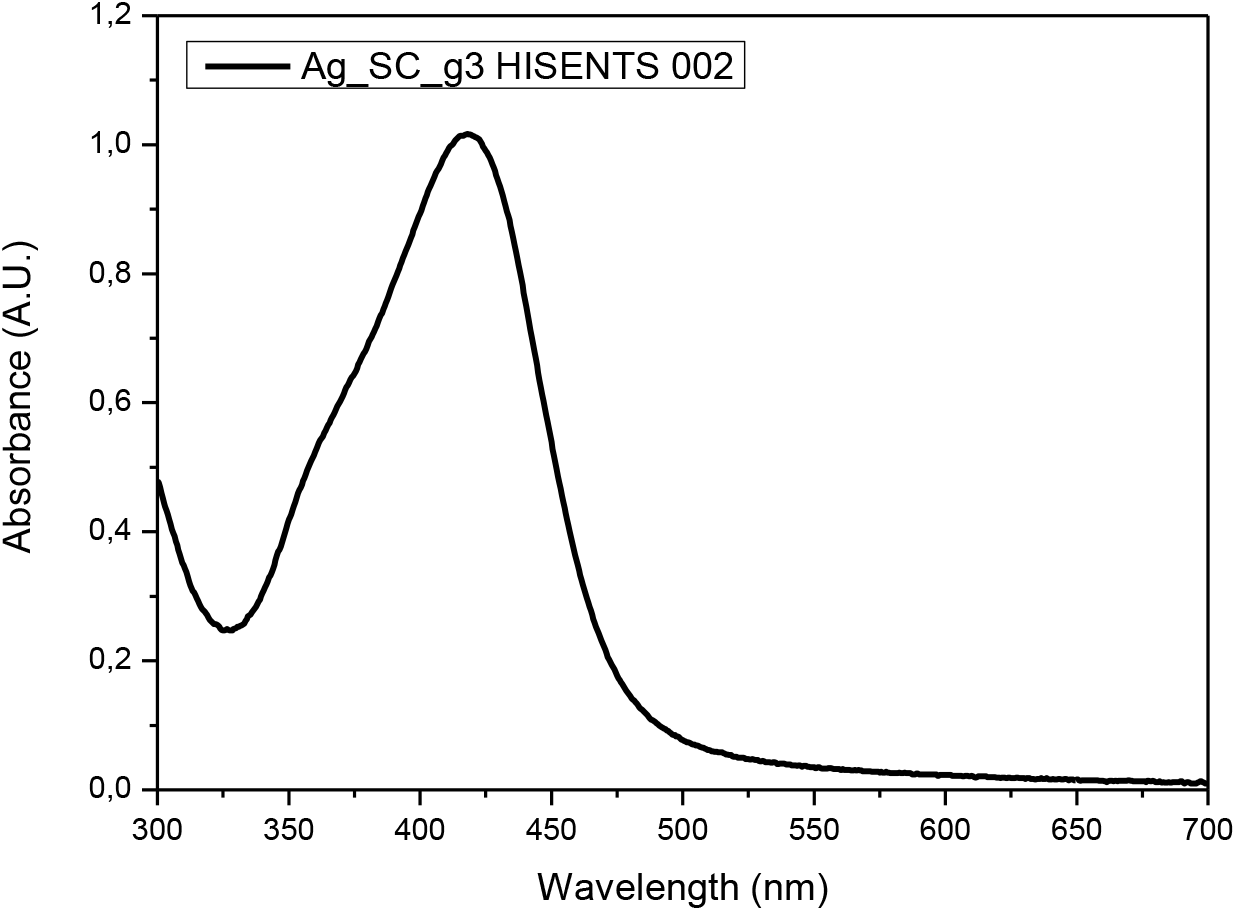
Experimental absobance spectrum of AgNPs. The maximum absobtion peak is observed at 417.4 nm.

### Leakage assay interference control

The dye 5(6)-Carboxyfluorescein (CF) was diluted in 20 mM HEPES 150 mM NaCl (HEPES saline buffer) or 20 mM HEPES 300mM glucose (HEPES glucose buffer) at concentrations from from 3×10^−4^ μM to 0.01 μM. Two sets of samples were prepared, one of them with 100 μM AgNPs and one without NPs. The samples were incubated 30 minutes and then the fluorescence intensity was measured at 514 nm.

**Figure S2.**
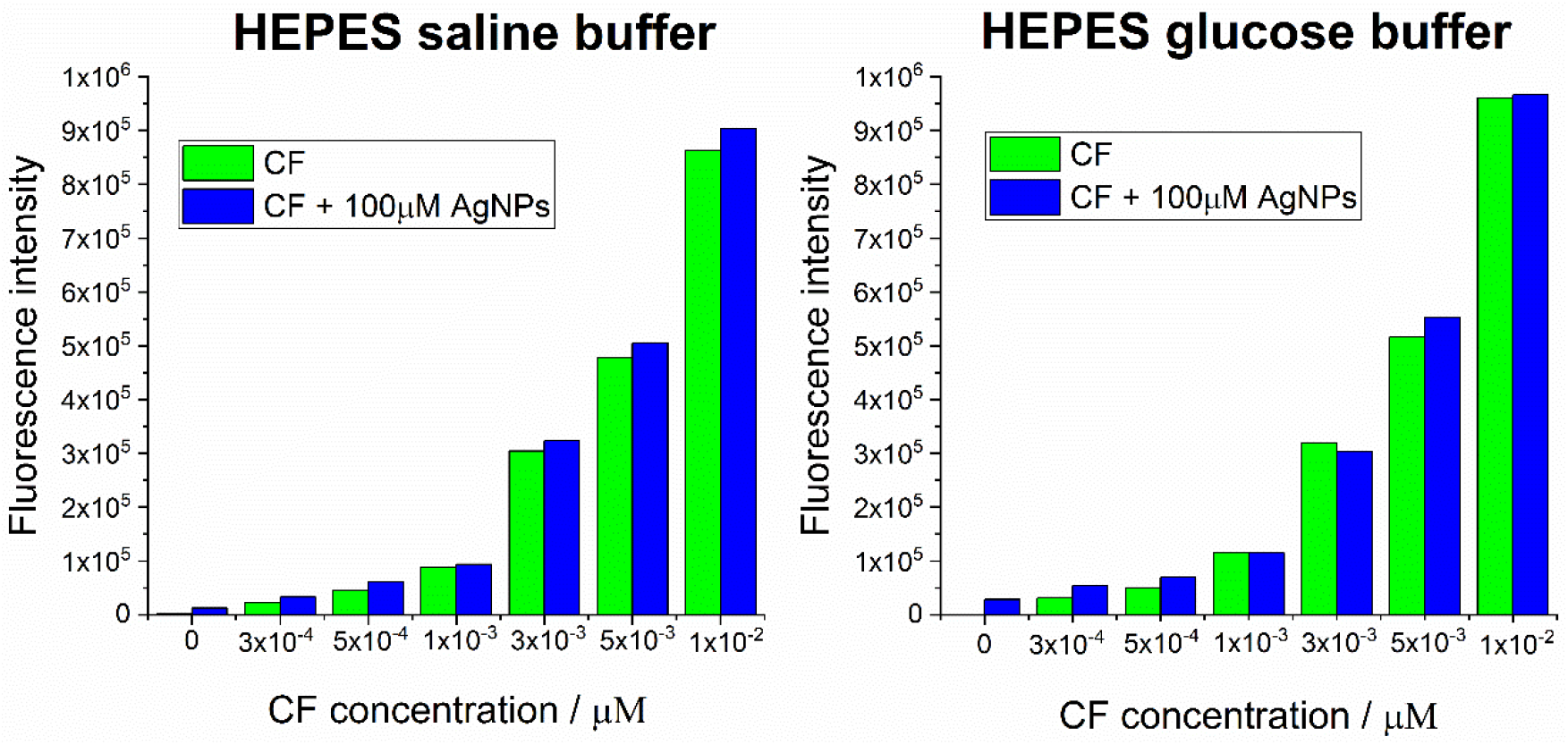
Comparison of fluorescence intensity signal of CF samples at different concentrations in the presence and abesence of 100 μM AgNPs. The presence of AgNPs barely affect the fluorescence intensity of CF.

## REFERENCES

(1) Srivastava, V.; Gusain, D.; Sharma, Y. C. Industrial & Engineering Chemistry Research 2015, 54, 6209.

(2) Zhang, L.; Gu, F. X.; Chan, J. M.; Wang, A. Z.; Langer, R. S.; Farokhzad, O. C. Clinical Pharmacology & Therapeutics 2008, 83, 761.

(3) Rascol, E.; Devoisselle, J. M.; Chopineau, J. Nanoscale 2016, 8, 4780.

(4) Singla, R.; Sharma, C.; Shukla, A. K.; Acharya, A. Journal of Nanoscience and Nanotechnology 2019, 19, 1889.

(5) Barbalinardo, M.; Caicci, F.; Cavallini, M.; Gentili, D. Small 2018, 14.

(6) Dillip, G. R.; Banerjee, A. N.; Sreekanth, T. V. M.; Anitha, V. C.; Joo, S. W. Materials Science in Semiconductor Processing 2017, 59, 87.

(7) Kim, I. Y.; Joachim, E.; Choi, H.; Kim, K. Nanomedicine-Nanotechnology Biology and Medicine 2015, 11, 1407.

(8) Calderon-Jimenez, B.; Johnson, M. E.; Bustos, A. R. M.; Murphy, K. E.; Winchester, M. R.; Baudrit, J. R. V. Frontiers in Chemistry 2017, 5.

(9) Vance, M. E.; Kuiken, T.; Vejerano, E. P.; McGinnis, S. P.; Hochella, M. F.; Rejeski, D.; Hull, M. S. Beilstein Journal of Nanotechnology 2015, 6, 1769.

(10) Sotiriou, G. A.; Pratsinis, S. E. Current Opinion in Chemical Engineering 2011, 1, 3.

(11) Piella, J.; Bastus, N. G.; Puntes, V. Zeitschrift Fur Physikalische Chemie-Internatonal Journal of Research in Physical Chemistry & Chemical Physics 2017, 231, 33.

(12) Li, Y. N.; Chang, Y. T.; Lian, X. F.; Zhou, L. Q.; Yu, Z. Q.; Wang, H. X.; An, F. F. Journal of Biomedical Nanotechnology 2018, 14, 1515.

(13) Dubey, P.; Matai, I.; Kumar, S. U.; Sachdev, A.; Bhushan, B.; Gopinath, P. Advances in Colloid and Interface Science 2015, 221, 4.

(14) Carlson, C.; Hussain, S. M.; Schrand, A. M.; Braydich-Stolle, L. K.; Hess, K. L.; Jones, R. L.; Schlager, J. J. Journal of Physical Chemistry B 2008, 112, 13608.

(15) Mukherjee, S. G.; O’Claonadh, N.; Casey, A.; Chambers, G. Toxicology in Vitro 2012, 26, 238.

(16) Xue, Y. Y.; Zhang, T.; Zhang, B. Y.; Gong, F.; Huang, Y. M.; Tang, M. Journal of Applied Toxicology 2016, 36, 352.

(17) Zou, Z. Z.; Chang, H. C.; Li, H. L.; Wang, S. M. Apoptosis 2017, 22, 1321.

(18) Volker, C.; Kampken, I.; Boedicker, C.; Oehlmann, J.; Oetken, M. Nanotoxicobgy 2015, 9, 677.

(19) Ahmed, K. B. R.; Nagy, A. M.; Brown, R. P.; Zhang, Q.; Malghan, S. G.; Goering, P. L. Toxicology in Vitro 2017, 38, 179.

(20) Casals, E.; Gonzalez, E.; Puntes, V. F. Journal of Physics D-Applied Physics 2012, 45.

(21) Nel, A. E.; Madler, L.; Velegol, D.; Xia, T.; Hoek, E. M. V.; Somasundaran, P.; Klaessig, F.; Castranova, V.; Thompson, M. Nature Materials 2009, 8, 543.

(22) Zhang, S. W.; Nelson, A.; Beales, P. A. Langmuir 2012, 28, 12831.

(23) Chithrani, B. D.; Ghazani, A. A.; Chan, W. C. W. Nano Letters 2006, 6, 662.

(24) Moghadam, B. Y.; Hou, W. C.; Corredor, C.; Westerhoff, P.; Posner, J. D. Langmuir 2012, 28, 16318.

(25) Montis, C.; Generini, V.; Boccalini, G.; Bergese, P.; Bani, D.; Berti, D. Journal of Colloid and Interface Science 2018, 516, 284.

(26) Werner, M.; Auth, T.; Beales, P. A.; Fleury, J. B.; Hook, F.; Kress, H.; Van Lehn, R. C.; Muller, M.; Petrov, E. P.; Sarkisov, L.; Sommer, J. U.; Baulin, V. A. Biointerphases 2018, 13.

(27) Behzadi, S.; Serpooshan, V.; Tao, W.; Hamaly, M. A.; Alkawareek, M. Y.; Dreaden, E. C.; Brown, D.; Alkilany, A. M.; Farokhzad, O. C.; Mahmoudi, M. Chemical Society Reviews 2017, 46, 4218.

(28) Beales, P. A.; Ciani, B.; Cleasby, A. J. Physical Chemistry Chemical Physics 2015, 17, 15489.

(29) Le, M. T.; Litzenberger, J. K.; Prenner, E. J. In Advances in Biomimetics; InTech: 2011.

(30) Bastus, N. G.; Merkoci, F.; Piella, J.; Puntes, V. Chemistry of Materials 2014, 26, 2836.

(31) Rouser, G.; Fleischer, S.; Yamamoto, A. Lipids 1970, 5, 494.

(32) Angelova, M. I.; Soleau, S.; Meleard, P.; Faucon, J. F.; Bothorel, P. Trends in Colloid and Interface Science VI 1992, 89, 127.

(33) Meleard, P.; Bagatolli, L. A.; Pott, T. Methods in Enzymology Liposomes, Pt G 2009, 465, 161.

(34) Pincet, F.; Adrien, V.; Yang, R.; Delacotte, J.; Rothman, J. E.; Urbach, W.; Tareste, D. Plos One 2016, 11.

(35) Soumpasis, D. M. Biophysical Journal 1983, 41, 95.

(36) Axelrod, D.; Koppel, D. E.; Schlessinger, J.; Elson, E.; Webb, W. W. Biophysical Journal 1976, 16, 1055.

(37) Rautu, S. A.; Orsi, D.; Di Michele, L.; Rowlands, G.; Cicuta, P.; Turner, M. S. Soft Matter 2017, 13, 3480.

(38) Bassereau, P.; Sorre, B.; Levy, A. Advances in Colloid and Interface Science 2014, 208, 47.

(39) Pecreaux, J.; Dobereiner, H. G.; Prost, J.; Joanny, J. F.; Bassereau, P. European Physical Journal E 2004, 13, 277.

(40) Bastus, N. G.; Casals, E.; Vazquez-Campos, S.; Puntes, V. Nanotoxicology 2008, 2, 99.

(41) Diegoli, S.; Manciulea, A. L.; Begum, S.; Jones, I. P.; Lead, J. R.; Preece, J. A. Science of the Total Environment 2008, 402, 51.

(42) Cumberland, S. A.; Lead, J. R. Journal of Chromatography A 2009, 1216, 9099.

(43) Huynh, K. A.; Chen, K. L. Environmental Science & Technology 2011, 45, 5564.

(44) Prathna, T. C.; Chandrasekaran, N.; Mukherjee, A. Colloids and Surfaces a- Physicochemical and Engineering Aspects 2011, 390, 216.

(45) Jang, M. H.; Lee, S.; Hwang, Y. S. Plos One 2015, 10.

(46) Li, S.; Malmstadt, N. Soft Matter 2013, 9, 4969.

(47) Leite, N. B.; Aufderhorst-Roberts, A.; Palma, M. S.; Connell, S. D.; Neto, J. R.; Beales, P. A. Biophysical Journal 2015, 109, 936.

(48) Bergstrom, C. L.; Beales, P. A.; Lv, Y.; Vanderlick, T. K.; Groves, J. T. Proceedings of the National Academy of Sciences of the United States of America 2013, 110, 6269.

(49) Yu, Y.; Granick, S. Journal of the American Chemical Society 2009, 131, 14158.

(50) Churchman, A. H.; Wallace, R.; Milne, S. J.; Brown, A. P.; Brydson, R.; Beales, P. A. Chemical Communications 2013, 49, 4172.

(51) Reynwar, B. J.; Illya, G.; Harmandaris, V. A.; Muller, M. M.; Kremer, K.; Deserno, M. Nature 2007, 447, 461.

(52) Spangler, E. J.; Kumar, P. B. S.; Laradji, M. Soft Matter 2018, 14, 5019.

(53) Saric, A.; Cacciuto, A. Physical Review Letters 2012, 108.

(54) Saric, A.; Cacciuto, A. Physical Review Letters 2012, 109.

(55) van den Bogaart, G.; Hermans, N.; Krasnikov, V.; de Vries, A. H.; Poolman, B. Biophysical Journal 2007, 92, 1598.

(56) Amjad, O. A.; Mognetti, B. M.; Cicuta, P.; Di Michele, L. Langmuir 2017, 33, 1139.

(57) Booth, A.; Marklew, C. J.; Ciani, B.; Beales, P. A. iScience 2019. DOI: 10.1016/j.isci.2019.04.021.

(58) Karatekin, E.; Sandre, O.; Guitouni, H.; Borghi, N.; Puech, P. H.; Brochard-Wyart, F. Biophysical Journal 2003, 84, 1734.

(59) Akimov, S. A.; Volynsky, P. E.; Galimzyanov, T. R.; Kuzmin, P. I.; Pavlov, K. V.; Batishchev, O. V. Scientific Reports 2017, 7.

(60) Fuertes, G.; Garcia-Saez, A. J.; Esteban-Martin, S.; Gimenez, D.; Sanchez-Munoz, O. L.; Schwille, P.; Salgado, J. Biophysical Journal 2010, 99, 2917.

(61) Delay, M.; Dolt, T.; Woellhaf, A.; Sembritzki, R.; Frimmel, F. H. Journal of Chromatography A 2011, 1218, 4206.

(62) Velegol, D. Journal of Nanophotonics 2007, 1.

(63) Montis, C.; Maiolo, D.; Alessandri, I.; Bergese, P.; Berti, D. Nanoscale 2014, 6, 6452.

(64) Lee, Y. K.; Choi, E. J.; Webster, T. J.; Kim, S. H.; Khang, D. International Journal of Nanomedicine 2015, 10, 97.

(65) Wei, X.; Qu, X.; Ding, L.; Hu, J.; Jiang, W. Environmental Pollution 2016, 219, 1.

(66) Di Silvio, D.; Maccarini, M.; Parker, R.; Mackie, A.; Fragneto, G.; Bombelli, F. B. Journal of Colloid and Interface Science 2017, 504, 741.

(67) Anderson, C. R.; Gnopo, Y. D. M.; Gambinossi, F.; Mylon, S. E.; Ferri, J. K. Journal of Biomedical Materials Research Part A 2018, 106, 1061.

(68) Lesniak, A.; Salvati, A.; Santos-Martinez, M. J.; Radomski, M. W.; Dawson, K. A.; Aberg, C. Journal of the American Chemical Society 2013, 135, 1438.

(69) Yi, P.; Chen, K. L. Environmental Science & Technology 2013, 47, 5711.

(70) Wang, Q. Y.; Lim, M. H.; Liu, X. T.; Wang, Z. W.; Chen, K. L. Environmental Science & Technology 2016, 50, 2301.

(71) Romer, W.; Berland, L.; Chambon, V.; Gaus, K.; Windschiegl, B.; Tenza, D.; Aly, M. R. E.; Fraisier, V.; Florent, J. C.; Perrais, D.; Lamaze, C.; Raposo, G.; Steinem, C.; Sens, P.; Bassereau, P.; Johannes, L. Nature 2007, 450, 670.

(72) Praper, T.; Sonnen, A. F. P.; Kladnik, A.; Andrighetti, A. O.; Viero, G.; Morris, K. J.; Volpi, E.; Lunelli, L.; Serra, M. D.; Froelich, C. J.; Gilbert, R. J. C.; Anderluh, G. Proceedings of the National Academy of Sciences of the United States of America 2011, 108, 21016.

(73) Solon, J.; Gareil, O.; Bassereau, P.; Gaudin, Y. Journal of General Virology 2005, 86, 3357.

(74) Lamaziere, A.; Burlina, F.; Wolf, C.; Chassaing, G.; Trugnan, G.; Ayala-Sanmartin, J. Plos One 2007, 2.

(75) Sandre, O.; Moreaux, L.; Brochard-Wyart, F. Proceedings of the National Academy of Sciences of the United States of America 1999, 96, 10591.

(76) Jaalouk, D. E.; Lammerding, J. Nature Reviews Molecular Cell Biology 2009, 10, 63.

(77) Gerhold, K. A.; Schwartz, M. A. Physiology 2016, 31, 359.

(78) Yasuda, N.; Miura, S. I.; Akazawa, H.; Tanaka, T.; Qin, Y.; Kiya, Y.; Imaizumi, S.; Fujino, M.; Ito, K.; Zou, Y.; Fukuhara, S.; Kunimoto, S.; Fukuzaki, K.; Sato, T.; Ge, J. B.; Mochizuki, N.; Nakaya, H.; Saku, K.; Komuro, I. Embo Reports 2008, 9, 179.

(79) Soubias, O.; Teague, W. E.; Hines, K. G.; Gawrisch, K. Biochimie 2014, 107, 28.

(80) Zhang, Y. L.; Frangos, J. A.; Chachisvilis, M. American Journal of Physiology-Cell Physiology 2009, 296, C1391.

(81) Chen, K. D.; Li, Y. S.; Kim, M.; Li, S.; Yuan, S.; Chien, S.; Shyy, J. Y. J. Journal of Biological Chemistry 1999, 274, 18393.

(82) Hoffman, B. D.; Grashoff, C.; Schwartz, M. A. Nature 2011, 475, 316.

(83) De Pascalis, C.; Etienne-Manneville, S. Molecular Biology of the Cell 2017, 28, 1833.

(84) Yamamoto, K.; Ando, J. Circulation Journal 2018, 82, 2691.

(85) Halder, G.; Dupont, S.; Piccolo, S. Nature Reviews Molecular Cell Biology 2012, 13, 591.

(86) Low, B. C.; Pan, C. Q.; Shivashankar, G. V.; Bershadsky, A.; Sudol, M.; Sheetz, M. Febs Letters 2014, 588, 2663.

(87) Mellouli, S.; Monterroso, B.; Vutukuri, H. R.; te Brinke, E.; Chokkalingam, V.; Rivas, G.; Huck, W. T. S. Soft Matter 2013, 9, 10493.

(88) Wei, Y.; Mayoral-Delgado, I.; Stewart, N. A.; Dymond, M. K. Chemistry and Physics of Lipids 2019, 218, 91.

